# Roq1 confers resistance to Xanthomonas, *Pseudomonas syringae* and *Ralstonia solanacearum* in tomato

**DOI:** 10.1101/813758

**Authors:** Nicholas C. Thomas, Connor G. Hendrich, Upinder S. Gill, Caitilyn Allen, Samuel F. Hutton, Alex Schultink

## Abstract

*Xanthomonas* species, *Pseudomonas syringae* and *Ralstonia solanacearum* are bacterial plant pathogens that cause significant yield loss in many crop species. Current control methods for these pathogens are insufficient but there is significant potential for generating new disease-resistant crop varieties. Plant immune receptors encoded by nucleotide-binding, leucine-rich repeat (NLR) genes typically confer resistance to pathogens that produce a cognate elicitor, often an effector protein secreted by the pathogen to promote virulence. The diverse sequence and presence / absence variation of pathogen effector proteins within and between pathogen species usually limits the utility of a single NLR gene to protecting a plant from a single pathogen species or particular strains. The NLR protein Recognition of XopQ 1 (Roq1) was recently identified from the plant *Nicotiana benthamiana* and mediates perception of the effector proteins XopQ and HopQ1 from *Xanthomonas* and *P. syringae* respectively. Unlike most recognized effectors, alleles of XopQ/HopQ1 are highly conserved and present in most plant pathogenic strains of *Xanthomonas* and *P. syringae*. A homolog of XopQ/HopQ1, named RipB, is present in many *R. solanacearum* strains. We found that Roq1 also mediates perception of RipB and confers immunity to *Xanthomonas, P. syringae*, and *R. solanacearum* when expressed in tomato. Strong resistance to *Xanthomonas perforans* was observed in three seasons of field trials with both natural and artificial inoculation. The *Roq1* gene can therefore be used to provide safe, economical and effective control of these pathogens in tomato and other crop species and reduce or eliminate the need for traditional chemical controls.

**Summary:** A single immune receptor expressed in tomato confers strong resistance to three different bacterial diseases.

## Introduction

Bacterial pathogens from the species *Pseudomonas syringae, Ralstonia solanacearum*, and the genus *Xanthomonas* can infect many different crop species and inflict significant yield losses when environmental conditions favor disease. *Xanthomonas* and *P. syringae* tend to enter plant stem, leaf, or flower tissue through wounds or natural openings, such as stomata or hydathodes, whereas *R. solanacearum* is soilborne, entering roots through wounds and natural openings before colonizing xylem tissue (Vasse, Frey, and Trigalet 1995; Gudesblat, Torres, and Vojnov 2009). Once inside the host these bacteria manipulate host metabolism and suppress plant immunity using multiple strategies, including effector proteins delivered by the type III secretion system (Kay and Bonas 2009; Peeters et al. 2013; Xin, Kvitko, and He 2018). This enables the pathogens to multiply to high titers while the plant tissue is still alive and showing few or no visual symptoms. Once the bacteria reach high populations they typically cause necrosis of infected leaf tissue or wilting and eventual death of the plant.

Effective control measures for bacterial pathogens are relatively limited, particularly once plants become infected (Davis et al. 2013). Soil fumigation can reduce *R. solanacearum* populations in the soil but this is expensive, potentially hazardous to workers and the environment, and of limited efficacy (Yuliar, Nion, and Toyota 2015). Copper sulfate and antibiotics such as streptomycin have been used to control *Xanthomonas* species and *P. syringae* but have adverse environmental impacts and many strains have evolved tolerance to these chemicals (Kennelly et al. 2007; Griffin et al. 2017). Applying chemicals that induce systemic acquired resistance, such as acibenzolar-S-methyl, can provide partial control but increase production cost and can depress crop yields when used repeatedly (Pontes et al. 2016).

The most effective, economical, and safe way to control bacterial pathogens is to plant crop varieties that are immune to the target pathogen (Jones et al. 2014; Vincelli 2016). Such immunity is often mediated by plant immune receptor genes. Plants have large families of cell surface and intracellular immune receptor proteins that surveil for the presence of invading pathogens (Zipfel 2014; Jones, Vance, and Dangl 2016). Effector proteins delivered by the bacterial type III secretion system are common ligands for intracellular plant immune receptors encoded by intracellular nucleotide-binding domain and leucine-rich repeat containing (NLR) genes (X. Li, Kapos, and Zhang 2015; Jones, Vance, and Dangl 2016; Kapos, Devendrakumar, and Li 2019). While effector proteins contribute to virulence on a susceptible host, an immune response is activated in the plant if that plant has the cognate receptor to recognize the effector. NLR genes typically confer strong, dominant resistance to pathogens that deliver the cognate recognized effector protein (Jones and Dangl 2006; Boller and He 2009; Deslandes and Rivas 2012; X. Li, Kapos, and Zhang 2015). Disease-resistant plants can be generated by identifying the appropriate plant immune receptor genes and transferring them into the target crop species (Dangl, Horvath, and Staskawicz 2013).

We recently identified the *Nicotiana benthamiana* immune receptor gene named *Recognition of XopQ 1* (*Roq1*), which appears to be restricted to the genus *Nicotiana* and is required for resistance to *Xanthomonas spp.* and *P. syringae* (Schultink et al. 2017). *Roq1* is a Toll/Interleukin-1 Receptor (TIR) NLR immune receptor that mediates recognition of the *Xanthomonas* effector protein XopQ and the homologous effector HopQ1 from *P. syringae*. XopQ is present in most species and strains of *Xanthomonas* (Ryan et al. 2011) and HopQ1 is present in 62% (290 of 467) sequenced putative pathogenic *P. syringae* strains (Dillon et al. 2019). XopQ/HopQ1 has homology to nucleoside hydrolases and has been shown to enhance virulence on susceptible hosts (Ferrante and Scortichini 2009; W. Li et al. 2013), possibly by altering cytokinin levels or interfering with the activity of host 14-3-3 proteins (W. Li et al. 2013; Giska et al. 2013; Teper et al. 2014; Hann et al. 2014). The conservation of XopQ/HopQ1 and their importance in virulence suggests that *Roq1* has widespread potential to confer resistance to these pathogens in diverse crop species. Indeed, transient expression assays demonstrated that *Roq1* can recognize XopQ/HopQ1 alleles from *Xanthomonas* and *P. syringae* pathogens of tomato, pepper, rice, citrus, cassava, brassica, and bean (Schultink et al. 2017). However, it was not known if *Roq1* can confer disease resistance when expressed in a crop plant.

Tomato is one of the most important vegetable crops and is highly susceptible to several bacterial diseases. Bacterial spot, bacterial speck and bacterial wilt of tomato are caused by *Xanthomonas* species, *P. syringae* pv. *tomato* and *R. solanacearum*, respectively. These diseases are difficult to control, especially if the pathogens become established in a field and environmental conditions favor disease (Rivard et al. 2012; Potnis et al. 2015). Tomato breeding germplasm has only limited resistance against these diseases and in some cases linkage drag has complicated introgression of resistance genes from wild relatives (Sharma and Bhattarai 2019). *R. solanacearum* contains a homolog of XopQ/HopQ1 called RipB, suggesting that expressing *Roq1* in tomato could also confer resistance to bacterial wilt. Like XopQ/HopQ1 in *Xanthomonas* and *P. syringae*, RipB is highly conserved in most *R. solanacearum* isolates (Sabbagh et al. 2019). Here we present laboratory and field data showing that *Roq1* confers resistance against these three pathogens in tomato and that this resistance depends on the presence of the cognate pathogen effector.

## Results

### Tomatoes expressing *Roq1* are resistant to *Xanthomonas* and *P. syringae*

We generated homozygous tomato plants expressing the *Roq1* gene from *N. benthamiana* and tested them for resistance to *Xanthomonas* and *P. syringae* by measuring bacterial growth *in planta*. Population sizes of wild-type *X. perforans* strain 4B and *X. euvesicatoria* strain 85-10 were approximately 100-fold smaller in tomatoes expressing *Roq1* compared to wild-type tomatoes at six days post inoculation (Fig. 1). In contrast, XopQ deletion mutants multiplied equally well in leaves of both wild-type and *Roq1* tomato. Disease symptoms begin as small water-soaked lesions and progress to necrosis of infected tissue. Wild-type *X. perforans* and *X. euvesicatoria* caused severe disease symptoms on wild-type tomato plants but failed to cause visible symptoms on *Roq1* plants (Fig. 2). The XopQ mutants caused similar disease symptoms on both wild-type and *Roq1* tomato. Similar results were observed for *P. syringae* DC3000 and its HopQ1 mutant (Fig. 1, 2) and a Race 1 isolate of *P. syringae* pv. *tomato* (Supplementary Fig. 1).

**Figure 1.**
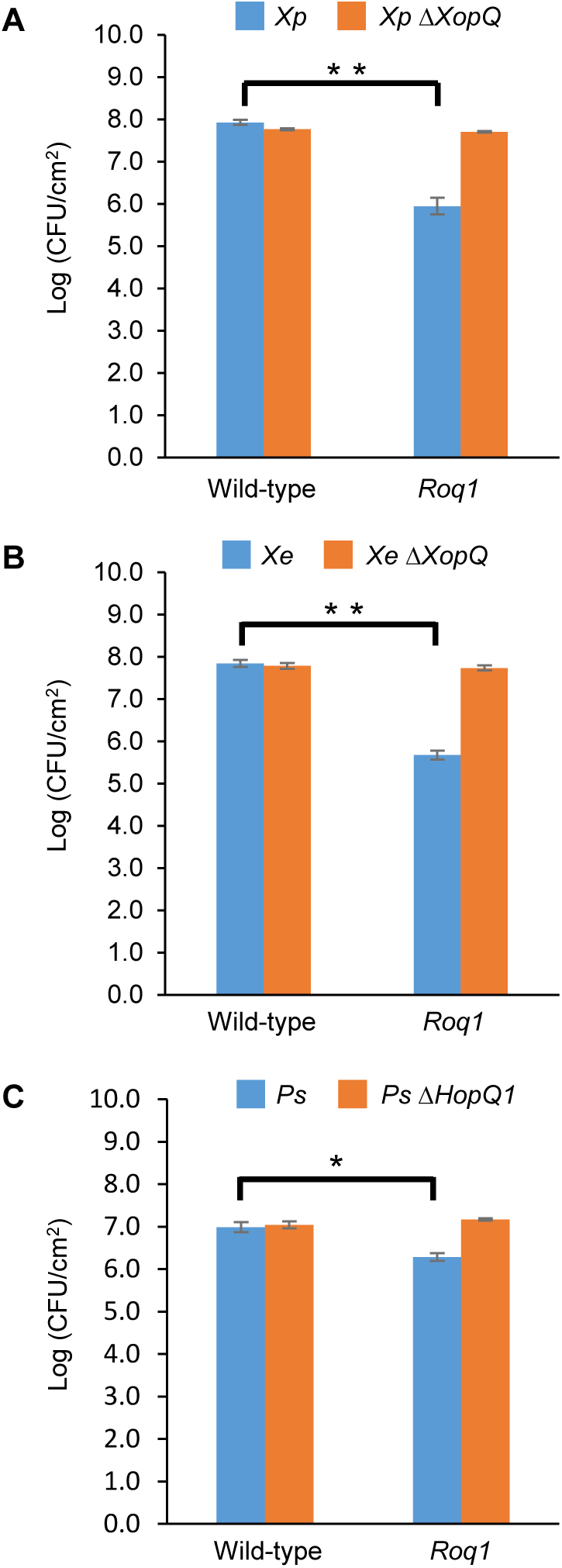
Bacterial growth in tomatoes expressing Roq1. *Xanthomonas perforans* 4B (*Xp*), *Xanthomonas euvesicatoria* 85-10 (*Xe*), and *Pseudomonas syringae* DC3000 (*Ps*) were infiltrated into leaf tissue of wild-type tomato and tomato expressing Roq1 at a low inoculum (OD_600_ = 0.0001 for Xe and Xp; OD_600_ = 0.00005 for Ps). Bacterial abundance was quantified by homogenizing leaf punches and counting colony forming units (CFU) per square centimeter of leaf tissue at six days post infiltration for *Xe* and *Xp;* three days post infiltration for *Ps*. Error bars indicate standard deviation. * = p < 0.05, ** = p < 0.01 by Student’s t-test.

**Figure 2.**
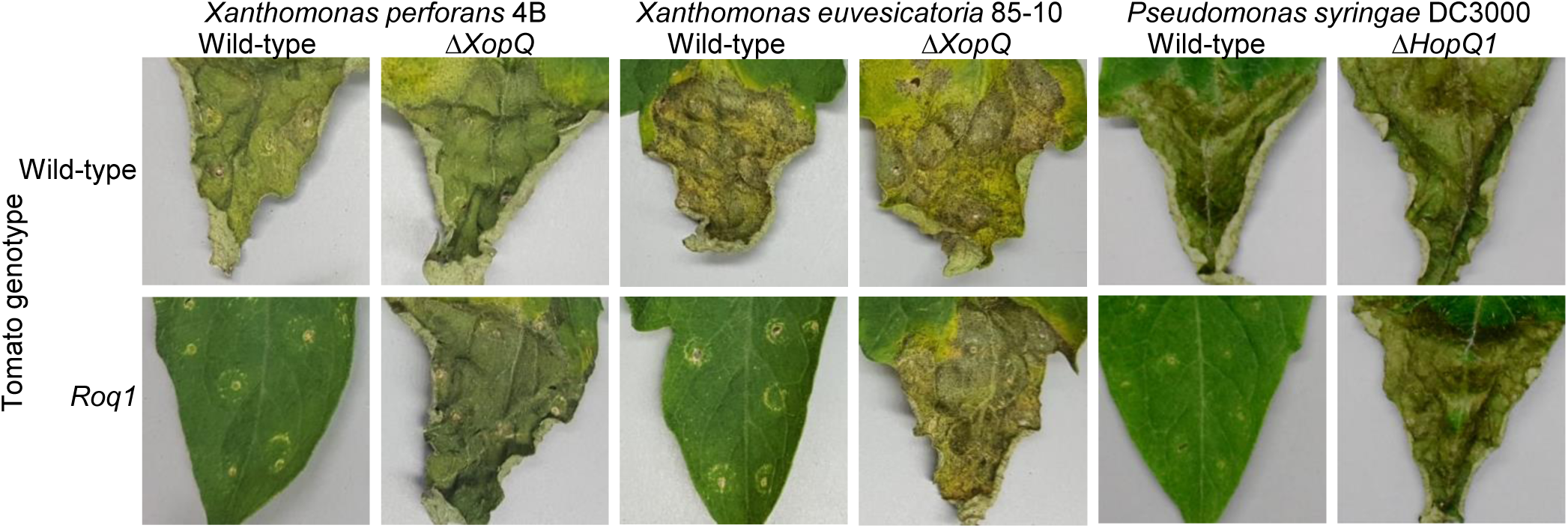
Bacterial disease symptoms on *Roq1* tomato. *Xanthomonas perforans* 4B (*Xp*), *Xanthomonas euvesicatoria* 85-10 (*Xe*), and *Pseudomonas syringae* DC3000 (*Ps*) wild-type and XopQ/HopQ1 knockout strains were infiltrated into tomato leaf tissue at low inoculum and disease symptoms were imaged at twelve, thirteen and four days post infiltration for *Xe, Xp* and *Ps* respectively. The infiltration was performed using a needless syringe and circular wounds from the infiltration are visible. The distal part of region of each leaf was infiltrated and the proximal part was left untreated. *Xe* and *Xp* were infiltrated at an OD_600_ of 0.0001 whereas *Ps* was infiltrated at an OD_600_ of 0.00005.

### Expression of *Roq1* confers resistance to *Xanthomonas perforans* in the field

To determine if the resistance observed in growth chamber experiments would hold up under commercial tomato production conditions, we tested the ability of *Roq1* tomatoes to resist *X. perforans* infection in the field. *Roq1* tomatoes were grown along with the Fla. 8000 wild-type parent as well as a Fla. 8000 variety expressing the Bs2 gene from pepper as a resistant control (Kunwar et al. 2018). For each of the three growing seasons both *Roq1* and the resistant *Bs2* control tomatoes showed significantly lower disease severity than the parental Fla. 8000 variety (Table 1) (p < 0.05). The total marketable yield of the *Roq1* plants was not significantly different from that of the susceptible parent for any of the three seasons (p > 0.05).

**Table 1.** Field trial results. A field trial was conducted in Florida with disease pressure from *Xanthomonas perforans*. Disease severity was scored out of 100 using the Horsfall-Barratt scale. Harvested tomatoes were graded and sized by USDA specifications to calculate the total marketable yield. The values shown are means ± standard deviation from at least four replicate plots of ten plants each. Tomato plants expressing the Bs2 immune receptor gene were included as a resistant control.

| Season /<br>Genotype | Disease<br>severity | Marketable yield<br>(kg/ha) |
| --- | --- | --- |
| **Spring 2018** |  |  |
| Fla. 8000 | 86 ± 5 | 54,655 ± 9,450 |
| Fla. 8000 Roq1 | 1 ± 1 | 52,656 ± 3,810 |
| Fla. 8000 Bs2 | 1 ± 1 | 66,270 ± 10,309 |
| **Fall 2018** |  |  |
| Fla. 8000 | 25 ± 7 | 19,576 ± 11,038 |
| Fla. 8000 Roq1 | 5 ± 1 | 18,538 ± 5,901 |
| Fla. 8000 Bs2 | 0 ± 0 | 33,770 ± 13,176 |
| **Spring 2019** |  |  |
| Fla. 8000 | 84 ± 2 | 73,009 ± 15,243 |
| Fla. 8000 Roq1 | 11 ± 7 | 92,837 ± 11,072 |
| Fla. 8000 Bs2 | 5 ± 1 | 80,516 ± 14,531 |

### The *R. solanacearum* RipB effector, a homolog of XopQ/HopQ1, is recognized by Roq1

RipB, considered a “core” effector of *R. solanacearum*, is present in approximately 90% of sequenced strains (Sabbagh et al. 2019), making it an attractive target ligand for engineering crop plants to be resistant to this pathogen. Roq1 perceives diverse alleles of XopQ and HopQ1 and we hypothesized that it can also recognize RipB. We constructed a phylogenetic tree of RipB, XopQ, and HopQ1 alleles identified by BLAST search and observed two major clades of RipB proteins, corresponding to phylotypes I and III (strains originating in Asia and Africa) and to phylotype II (strains originating in the Americas) (Fig. 3). We selected RipB alleles from *R. solanacearum* strains GMI1000 and MolK2, which are present in clades 1 and 2, respectively, for subsequent analysis. These two RipB alleles share 71% amino acid identity with each other and approximately 52% identity with XopQ excluding the divergent N terminus containing the putative type III secretion signal. An alignment of these two RipB proteins with XopQ and HopQ1 is shown in Supplementary Fig. 2. To test for Roq1-dependent recognition of RipB, we used *Agrobacterium* to transiently express RipB from GMI1000 and Molk2 in leaf tissue of wild-type and *roq1* mutant *N. tabacum*. Both RipB alleles triggered a strong hypersensitive / cell death response in wild-type *N. tabacum*, indicating immune activation. This response was absent in the *roq1-1* mutant but could be restored by transiently expressing Roq1 along with XopQ, RipB_GMI1000_, or RipB_Molk2_ (Fig. 4).

**Figure 3.**
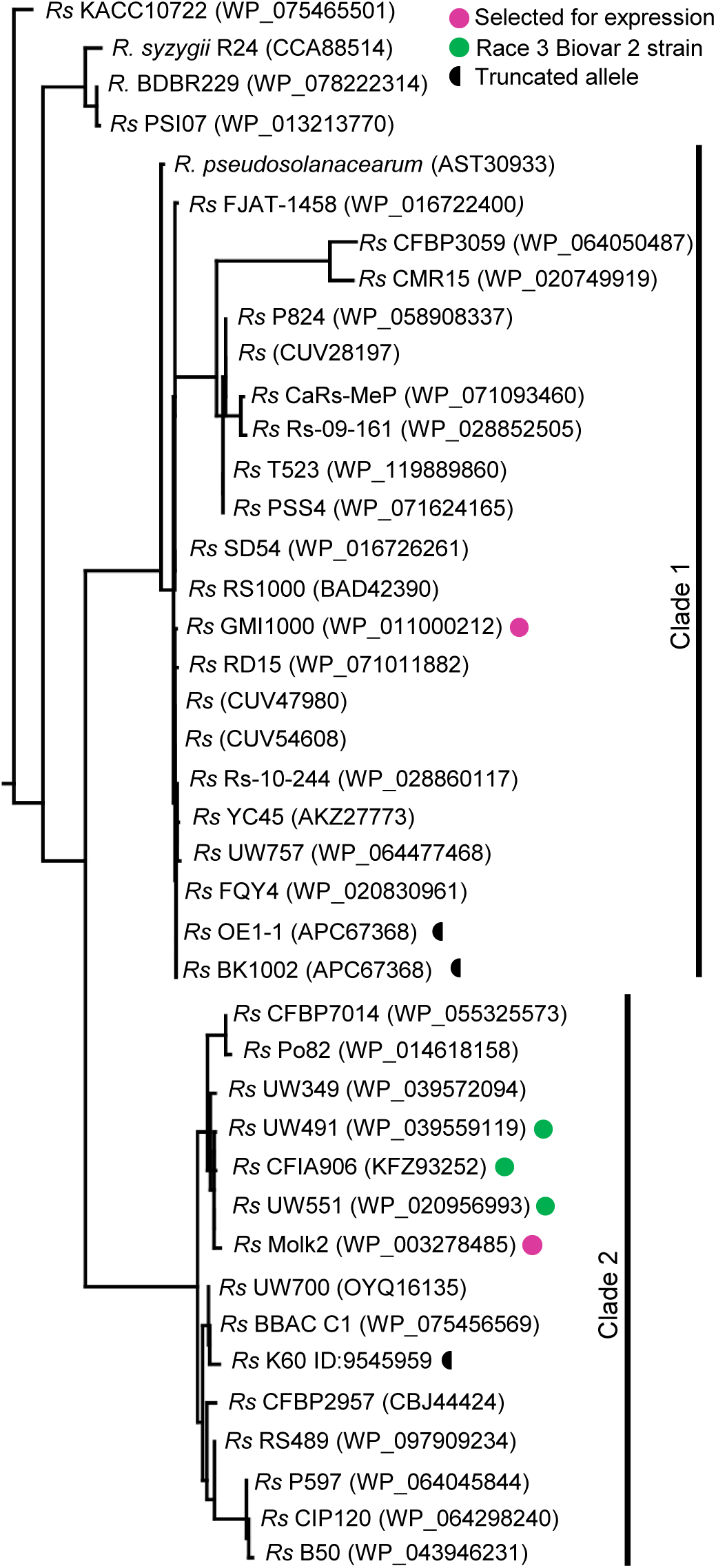
Phylogenetic tree of RipB proteins. Two major clades of RipB alleles from *Ralstonia solanacearum* strains are visible in a maximum likelihood tree generated from XopQ, HopQ1 and RipB protein sequences. RipB alleles from *Ralstonia solanacearum* strains GMI1000 and MolK2 were cloned for testing in this study and are indicated by pink dots. Several alleles have truncations that may make them nonfunctional (indicated by a half circle). Abbreviations were used for *Ralstonia* (R) *and Ralstonia solanacearum* (Rs).

**Figure 4.**
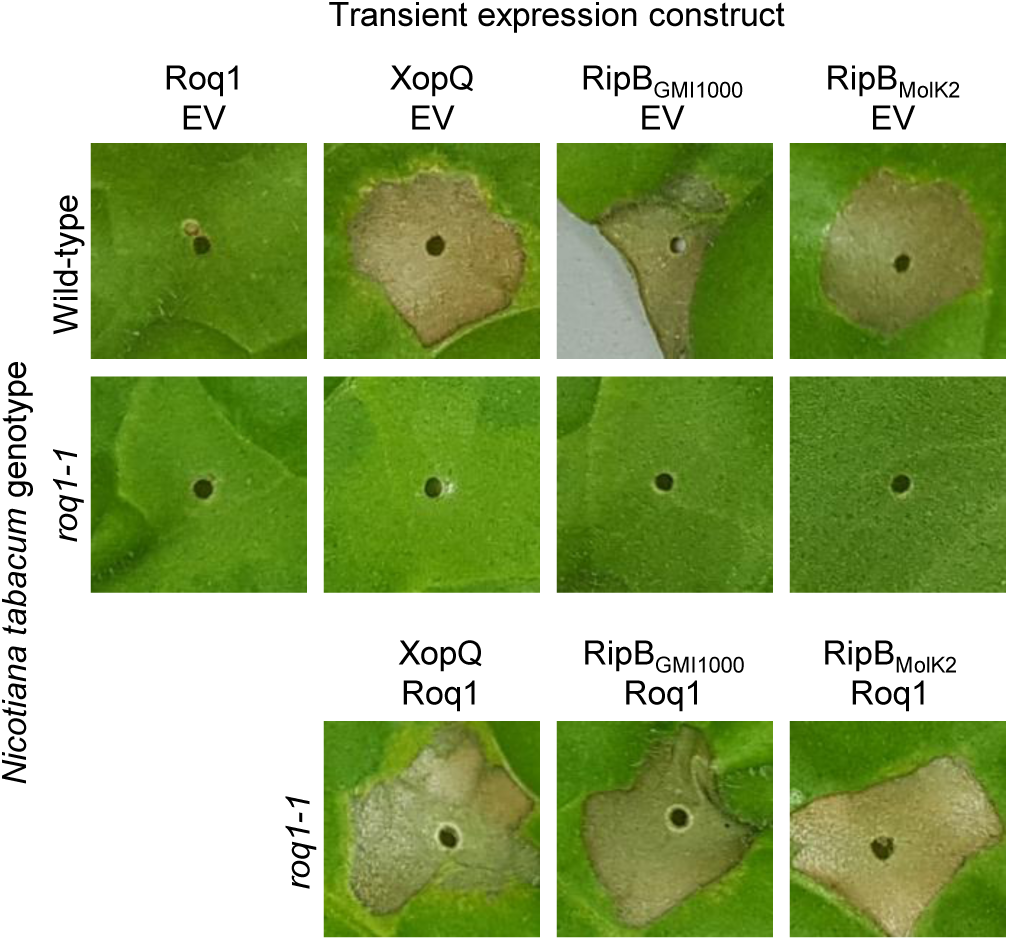
Roq1-dependent recognition of RipB in *Nicotiana tabacum. Agrobacterium tumefaciens* was used to transiently expressed XopQ, RipB_GMI1000_ and RipB_MolK2_ along with either Roq1 or an empty vector (EV) control in wild-type *Nicotiana tabacum* and a *roq1* loss of function mutant. The *Agrobacterium* was infiltrated at a total OD_600_ of 0.5 and the leaves were imaged at three days post infiltration.

### *Roq1* tomatoes are resistant to *R. solanacearum* containing RipB

Our observation that Roq1 can recognize RipB in leaf transient expression assays suggested that Roq1 can mediate resistance to bacterial wilt caused by *R. solanacearum*. We tested this hypothesis by challenging wild-type and Roq1-expressing tomato plants with *R. solanacearum* strain GMI1000 using a soil soak inoculation disease assay. Wild-type plants developed severe wilting approximately seven days after inoculation, whereas *Roq1* tomato plants remained mostly healthy over the two-week time course. The *Roq1* tomato plants were susceptible to a deletion mutant lacking RipB (GMI1000 *ΔripB*) (Fig. 5a). We also challenged plants by introducing bacteria directly to the xylem by placing bacteria on the surface of a cut petiole. Wild-type plants were wilted at eight days whereas *Roq1* plants remained healthy (Fig. 5b). Tomatoes expressing *Roq1* were also resistant to *R. solanacearum* strain UW551, which is a Race 3 Biovar 2 potato brown rot strain that has a clade 2 RipB allele (Supplementary Fig. 3).

**Figure 5.**
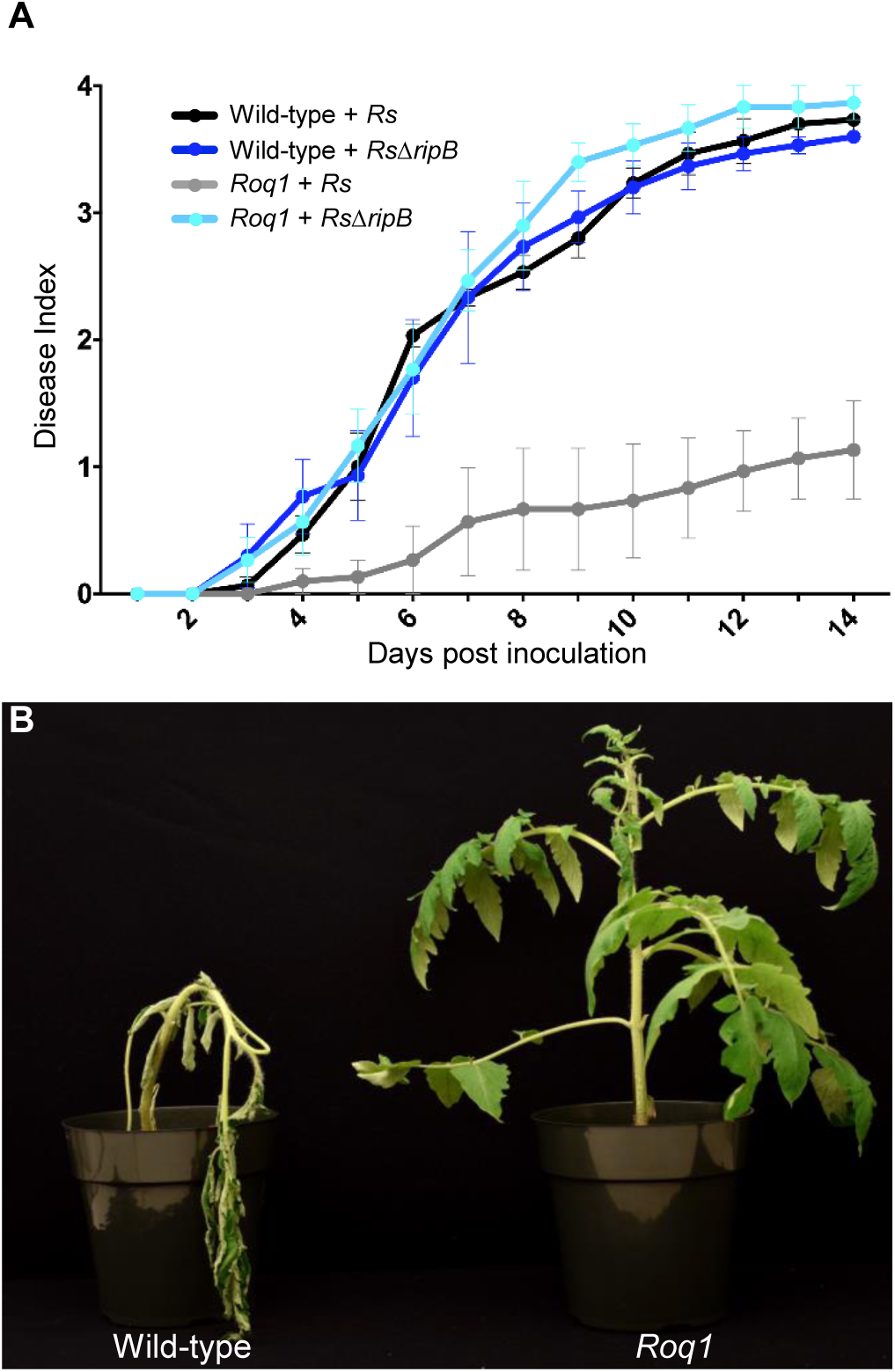
Bacterial Wilt disease development in Roq1 tomatoes. Wild-type and Roq1 tomatoes were infected with wild-type and RipB mutant *Ralstonia solanacearum* strain GMI1000 by soil soak inoculation. Disease symptoms were monitored over 14 days, with no wilting corresponding to a Disease Index of 0 and complete wilting corresponding to a Disease Index of 4. Error bars indicate standard error. (B) Wild-type and Roq1 tomato plants 8 days after petiole inoculation with approximately 2,000 cells of wild-type GMI1000.

### Occurrence of RipB in the *R. solanacearum* species complex

To investigate the potential for using *Roq1* to protect plants from *R. solanacearum*, we investigated the occurrence of RipB alleles in select *R. solanacearum* strains. Table 2 summarizes published known hosts of several *R. solanacearum* strains along with the phylotype and identified RipB allele. All strains in Table 2 except for tobacco pathogenic strains K60, Y45, BK1002 and OE1-1 contain putative full-length and functional RipB alleles. Relative to other RipB alleles, the K60 RipB allele is truncated at residue 437 and missing approximately 65 C-terminal residues and the OE1-1 allele is truncated at residue 425, missing approximately 77 residues based on a published genome sequence (Hayes, MacIntyre, and Allen 2017). Y45 does not have a predicted RipB allele based on a draft genome sequence (Z. Li et al. 2011).

**Table 2.** RipB occurrence and host range in *Ralstonia solanacearum*. The published host range of select *Ralstonia solanacearum* strains is listed along with the identified RipB allele. Truncated indicates that the identified allele is missing conserved residues at the C terminus and is putatively non-functional. * indicates a Race 3 Biovar 2 Select Agent strain.

| Strain | Host(s) | Phylotype | RipB allele | RipB accession |
| --- | --- | --- | --- | --- |
| GMI1000 | Tomato, Pepper, Arabidopsis | I | Present | WP_011000212 |
| RS1000 | Tomato | I | Present | BAD42390 |
| OE1-1 | Tobacco | I | Truncated | APC67368 |
| BK1002 | Tobacco |  | Truncated | LC459955 |
| Y45 | Tobacco | IB | Absent |  |
| K60 | Tomato, Tobacco | IIA | Truncated | CCF97494 |
| CFBP2957 | Tomato | IIA | Present | CBJ44424 |
| Po82 | Tomato, Banana, Potato | IIB | Present | WP_014618158 |
| IPO1609* | Potato | IIB | Present | WP_020956993 |
| MolK2 | Banana | IIB | Present | WP_003278485 |
| UW551* | Geranium, Tomato | IIB | Present | EAP72492 |
| CMR15 | Tomato | III | Present | WP_020749919 |
| PSI07 | Tomato | IV | Present | WP_013213770 |
| BDB R229 | Banana |  | Present | WP_078222314 |

## Discussion

*Roq1* expression in tomato confers strong resistance to *X. perforans, X. euvesicatoria and P. syringae* pv. *tomato*. Its effectiveness is dependent on the presence of the recognized effector protein XopQ/HopQ1 (Fig. 1, 2). Field trials revealed that tomatoes expressing *Roq1* were less susceptible to *X. perforans* than wild-type tomatoes in conditions approximating commercial production (Table 1). *Roq1* conferred a similar level of resistance as the Bs2-containing resistant check variety in one season and was slightly weaker in the other two. Bacterial spot caused by *X. perforans* can cause lesions on fruits, making them unsuitable for commercial sale, and also reduce plant productivity by damaging leaf tissue. Despite showing strong disease resistance, the tested *Roq1* line did not give a significantly greater total marketable yield than the susceptible parental variety. Of the three seasons, Spring 2019 had weather conditions expected to be most conducive for observing an impact of bacterial spot on marketable yield with mid-season rain promoting the early development of disease symptoms. The average marketable yield for the *Roq1* tomatoes was 27% higher than wild-type in this season, although a relatively small sample size (4 replicate plots of ten plants each) and a large variability of yield between plots resulted in a p-value of 0.08 by Student’s t-test. Although we cannot conclude that *Roq1* improves marketable yield of tomatoes from this data, a larger trial under high disease pressure may reveal such a difference.

It was unclear if *Roq1* could confer resistance to *R. solanacearum* because it colonizes different tissues than *Xanthomonas* and *P. syringae*. While *Xanthomonas* and *P. syringae* colonize tomato leaf tissue, *R. solanacearum* enters through the roots and colonizes xylem vessels. Although *R. solanacearum’s* type III secretion system is essential for virulence, it is not clear when and where the pathogen delivers effectors into host cells. It was therefore not clear if *Roq1* would be able to confer resistance to this pathogen in tomato. Here we confirmed that tomato plants expressing *Roq1* had strong resistance to *R. solanacearum* expressing RipB as measured by both soil soak and cut-petiole inoculation assays. This result is consistent with and expands on the recent report that RipB is recognized in *Nicotiana* species and that silencing *Roq1* confers susceptibility to *R. solanacearum* in *Nicotiana benthamiana(Nakano and Mukaihara 2019)*. In addition, *Roq1* confers resistance to *R. solanacearum* Race 3 biovar 3 strain UW551, a pathogen that can overcome other known sources of bacterial wilt resistance in tomato(Milling, Babujee, and Allen 2011). Some but not all of the *Roq1* tomatoes inoculated by the soil soak assay were colonized by *R. solanacearum* (Supplementary Fig. 4), implying that Roq1 both restricts the establishment of vascular colonization and separately reduces bacterial titers if colonization does occur. Activation of immune receptors, including Roq1, is known to induce many defense-associated genes with different putative activities (Sohn et al. 2014; Qi et al. 2018), presumably acting to inhibit pathogen virulence by distinct mechanisms. The observation that Roq1 inhibits both colonization establishment and population growth suggest that at least two independent downstream defense responses mediate the observed resistance phenotype.

The *Roq1* tomatoes were fully susceptible to an *R. solanacearum* mutant lacking *RipB*, indicating that the resistance depends on the interaction between RipB and Roq1. This is consistent with the observation that several naturally occurring *R. solanacearum* strains that can infect tobacco have truncated or are missing the RipB effector (Table 2)(Nakano and Mukaihara 2019), suggesting that losing RipB can allow the pathogen to overcome the native *Roq1* gene present in *Nicotiana tabacum*. Tobacco-infecting strains K60 and OE1-1 contain independently truncated RipB alleles (Fig. 3) and there have likely been multiple independent gene loss events which enable strains to evade Roq1-mediated resistance. Similarly, HopQ1 has been lost in strains of *P. syringae* that can infect tobacco (Denny 2006; Ferrante and Scortichini 2009; Z. Li et al. 2011). This suggests that this effector is not essential for virulence and it would therefore be prudent to deploy *Roq1* in combination with other disease resistance traits to avoid resistance breakdown due to pathogens losing XopQ/HopQ1/RipB.

No other known NLR immune receptor confers resistance against such a broad range of bacterial pathogens as Roq1. Effectors that are recognized by NLR proteins act as avirulence factors and are under strong evolutionary pressure to diversify or be lost to evade immune activation. Therefore the effector repertoires of pathogens are often quite diverse, with relatively few “core” effectors conserved within a species and even fewer shared between different genera (Grant et al. 2006). Effectors recognized by plant NLRs are typically narrowly conserved within a single bacterial genus (Kapos, Devendrakumar, and Li 2019). One such effector is AvrBs2, recognized by the Bs2 receptor from pepper, which is present in many *Xanthomonas* strains but is absent from *P. syringae* and *R. solanacearum*. In contrast XopQ/HopQ1/RipB is highly conserved in most *Xanthomonas, P. syringae* and *R. solanacearum* strains that cause disease in crop plants including kiwi (*P. syringae* pv. *actinidae*), banana (*R. solanacearum* and *X. campestris* pv. *musacearum*), stone fruit (*P. syringae*), pepper (*X. euvesicatoria*), citrus (*X. citri*), strawberry (*X. fragariae*), brassica (*X. campestris*), rice (*X. oryzae*), potato (*R. solanacearum*) and others. *R. solanacearum* Race 3 Biovar 2 strains are of particular concern because they are cold tolerant and potentially threaten potato cultivation in temperate climates. As a result, *R. solanacearum* Race 3 biovar 2 is a strictly regulated quarantine pathogen in Europe and North America and is on the United States Select Agent list. The ability of *Roq1* to protect tomato from the Race 3 Biovar 2 strain UW551 (Supplementary Fig. 3) suggests that *Roq1* can also protect potato from this high-concern pathogen. This work demonstrates the widespread potential of using naturally occurring plant immune receptors to manage diverse and difficult to control pathogen species safely, sustainably and economically.

## Methods

### Generation of tomato expressing Roq1

The Roq1 coding sequence was amplified from *N. benthamiana* cDNA and cloned into the pORE E4 binary plasmid (Coutu et al. 2007). *A. tumefaciens* co-cultivation was used to transform *Roq1* into the commercial tomato variety Fla. 8000 at the University of Nebraska Plant Transformation Core Research Facility. Transformed plants were selected by resistance to kanamycin, confirmed by genotyping, and selfed to obtain homozygous lines.

### Bacterial Leaf Spot and Leaf Speck disease assays

*Xanthomonas spp.* cultures were grown in NYG broth (0.5% peptone, 0.3% yeast extract, 2% glycerol) with rifampicin (100 μg / mL) overnight at 30 °C. *P. syringae* cultures were grown in KB broth (1% peptone, 0.15% K_2_HPO_4_, 1.5% glycerol, 5 mM MgSO_4_, pH 7.0) with rifampicin (100 μg / mL) overnight at 28 °C. Bacterial cultures were spun down at 5200 *g*, washed once with 10 mM MgCl_2_, and then diluted to the appropriate infiltration density with 10 mM MgCl_2_. Leaf tissue of tomato plants (approximately four weeks old) was infiltrated with bacterial solution using a needleless syringe. To quantify bacterial growth, leaf punches were homogenized in water, serially diluted and plated on NYG (for *Xanthomonas spp.*) or KB (for *P. syringae*) plates supplemented with 100 μg / mL rifampicin and 50 μg / mL cycloheximide to measure colony forming units. *X. perforans* strain 4B, *X. euvesicatoria* strain 85-10, and *P. syringae* strain DC3000 and the corresponding XopQ/HopQ1 deletion mutants were described previously (Schwartz et al. 2015; Schultink et al. 2017). The *P. syringae* pv. *tomato* Race 1 strain was isolated from a field of tomatoes with the PTO resistance gene in 1993 in California.

### Transient expression of RipB and XopQ

Alleles of RipB from *R. solanacearum* (NCBI Genbank accessions CAD13773.2 and WP_003278485) were synthesized and cloned into a BsaI-compatible version of the pORE E4 vector (Coutu et al. 2007). This plasmid was transformed into *A. tumefaciens* strain C58C1. *A. tumefaciens* cultures were grown on a shaker overnight at 30 °C in LB broth with rifampicin (100 μg / mL), tetracycline (10 μg / mL) and kanamycin (50 μg / mL). The cells were collected by centrifugation and resuspended in infiltration buffer (10 mM 2-(N-morpholino)ethanesulfonic acid, 10 mM MgCl_2_, pH 5.6), and diluted to an OD_600_ of 0.5 for infiltration into *N. tabacum* leaf tissue.

### *N. tabacum roq1* mutant lines

*N. tabacum roq1* mutant lines were generated by transforming *N. tabacum* with a construct coding for CAS9 and a guide RNA targeting the Roq1 gene with the sequence GATGATAAGGAGTTAAAGAG. This construct was also used for the generation of *N. benthamiana roq1* mutants published in Qi et al. 2018. Transformed *N. tabacum* plants were generated by Agrobacterium co-cultivation and selected for using kanamycin. Transformed plants were genotyped for the presence of mutations at the target site by PCR and Sanger sequencing. The *N. tabacum* mutant *roq1-1* has a single base pair A insertion at the target cut site in the *Roq1* gene.

### Bacterial wilt virulence assays

*R. solanacearum* virulence on tomato was measured as previously described(Khokhani et al. 2018). Briefly, cells of *R. solanacearum* strain GMI1000 grown overnight in CPG (0.1% casein hydrolysate, 1% peptone, 0.5% glucose, pH 7.0) at 28°C were collected by centrifugation and diluted to an OD_600_ of 0.1 in water (1×10^8^ CFU/ml). 50 mL of this suspension was poured on the soil around 17-day old tomato plants. Disease was rated daily for two weeks on a 0-4 disease index scale, where 0 is no leaves wilted, 1 is 1-25% wilted, 2 is 26-50% wilted, 3 is 51-75% of wilted, and 4 is 76-100% wilted. Data represent a total of four biological replicates with ten plants per replicate. Virulence data were analyzed using repeated measures ANOVA (Khokhani et al 2018). For petiole infection, the petiole of the first true leaf was cut with a razor blade horizontally approximately 1 cm from the stem. A drop of bacterial solution (2 uL, OD_600_ = 0.001) was pipetted onto the exposed cut petiole surface.

### Field trial disease assays

Three field trials were conducted at the University of Florida Gulf Coast Research and Education Center in Balm during the spring seasons of 2018 and 2019 and the fall season of 2018 and under the notification process of the United States Department of Agriculture. Large-fruited, fresh market tomato lines were used in these trials and included the inbred line, Fla. 8000, and nearly-isogenic lines containing either *Roq1* (event 316.4) or *Bs2* (Kunwar et al. 2018); along with commercial hybrids Florida 91, Sanibel, and HM 1823 as additional, susceptible controls (data not shown). For each trial, seeds were sown directly into peat-lite soilless media (Speedling, Sun City, FL, USA) in 128-cell trays (38 cm3 cell size). Transplants were grown in a greenhouse until 5 or 6 weeks, then planted to field beds that had been fumigated and covered with reflective plastic mulch. Field trials were conducted using a randomized complete block design with four blocks and 10-plant plots. Field plants were staked and tied, and irrigation was applied through drip tape beneath the plastic mulch of each bed. A recommended fertilizer and pesticide program were followed throughout the growing season, excluding the use of plant defense inducers, copper, or other bactericides (Freeman et al. 2018). Fruit were harvested from the inner 8 plants of each plot at the breaker stage and beyond graded for marketability according to USDA specifications. Yield data were analyzed using the PROC GLIMMIX procedure in SAS (version 9.4; SAS Institute, Cary, NC, USA), and block was considered random effects.

Field trials were inoculated with *X. perforans* race T4 (strain mixture of GEV904, GEV917, GEV1001, and GEV1063). Bacterial strains were grown on nutrient agar medium (BBL, Becton Dickinson and Co., Cockeysville, MD) and incubated at 28 °C for 24 h. Bacterial cells were removed from the plates and suspended in a 10 mM MgSo_4_·7H2O solution, and the suspension was adjusted to OD_600_=0.3, which corresponds to 10^8^ CFU/ml. The suspension for each strain was then diluted to 10^6^ CFU/ml, mixed in equal volume, and applied along with polyoxyethylene sorbitan monolaurate (Tween 20; 0.05% [vol/vol]) for field inoculation. Field trial plants were inoculated approximately 3 weeks after transplanting.

Bacterial spot disease severity was recorded three to eight weeks after inoculation using the Horsfall-Barratt scale (Horsfall, JG and Barrat, RW 1945), and ratings were converted to midpoint percentages for statistical analysis. Disease severity data were analyzed using a nonparametric procedure for the analysis of ordinal data (Brunner and Puri 2001; Shah and Madden 2004). Analysis of variance type statistic of ranked data was conducted using the PROC MIXED procedure in SAS. Relative marginal effects (RME) were generated with the equation: RME = (*R* – 0.5)/*N*; where *R* is the mean treatment ranking, and *N* is the total number of experimental units in the analysis; the LD_CI macro was used to generate 95% confidence intervals (Brunner and Puri 2001; Shah and Madden 2004). Blocks were considered random effects.

### Generation of the *R. solanacearum ΔripB* mutant

An unmarked *ΔripB* mutant was created using *sacB* positive selection with the vector pUFR80 (Castañeda et al. 2005). Briefly, the regions upstream and downstream of *ripB* were amplified using the primers ripBupF/R and ripBdwnF/R. These fragments were inserted into pUFR80 digested with HindIII and EcoRI using Gibson Assembly (Gibson et al. 2009) (New England Biolabs, Ipswitch, MA) and this construct was incorporated into the genome of strain GMI1000 using natural transformation, with successful integrants selected on CPG + kanamycin (Coupat et al. 2008). Plasmid loss was then selected for on CPG plates containing 5% w/v sucrose.Correct deletions were confirmed using PCR and sequencing.

### Phylogenetic analysis of XopQ, HopQ1 and RipB alleles

RipB alleles were identified by BLAST search of the NCBI protein database. Clustal Omega (Sievers et al. 2011) was used to generate a multiple sequence alignment with XopQ and HopQ1 alleles. To span the diversity of RipB alleles without have many redundant sequences, only a single sequence was retained if there were multiple identical or nearly identical sequences identified. A maximum likelihood tree was generated using PhyML (Guindon et al. 2010).

## Acknowledgements

We thank Shirley Sato and Tom Clemente of the University of Nebraska Plant Transformation Core Research Facility for the transformation of tomato. We thank Myeong-Je Cho and Julie Pham of the UC Berkeley Innovative Genomics Institute for transformation of *Nicotiana tabacum*.

