## Supplemental Data for "Roq1 confers resistance to Xanthomonas, *Pseudomonas syringae* and *Ralstonia solanacearum* in tomato"

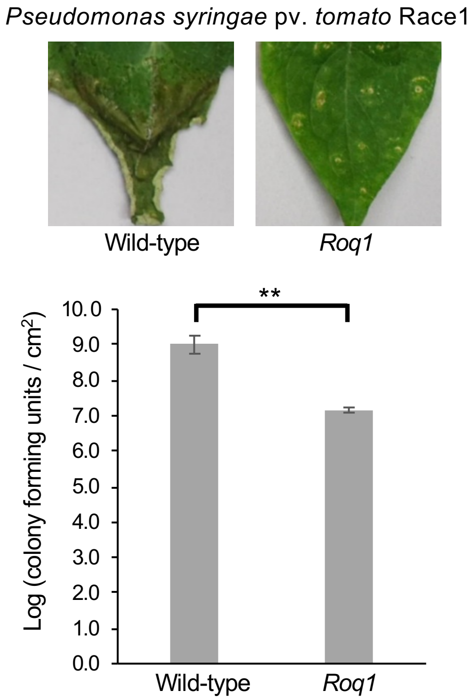


**Supplementary Figure 1.** Growth of *Pseudomonas* Race 1 in wild-type and *Roq1* tomatoes. *Pseudomonas syringae* pv. *tomato* Race 1 was infiltrated into wild-type and Roq1 tomatoes. At four days post infiltration disease symptoms were imaged (top) and colony forming units were determined by dilution plating of homogenized tissue. Error bars indicate standard deviation. ** = $p < 0.01$ by Student's t-test.

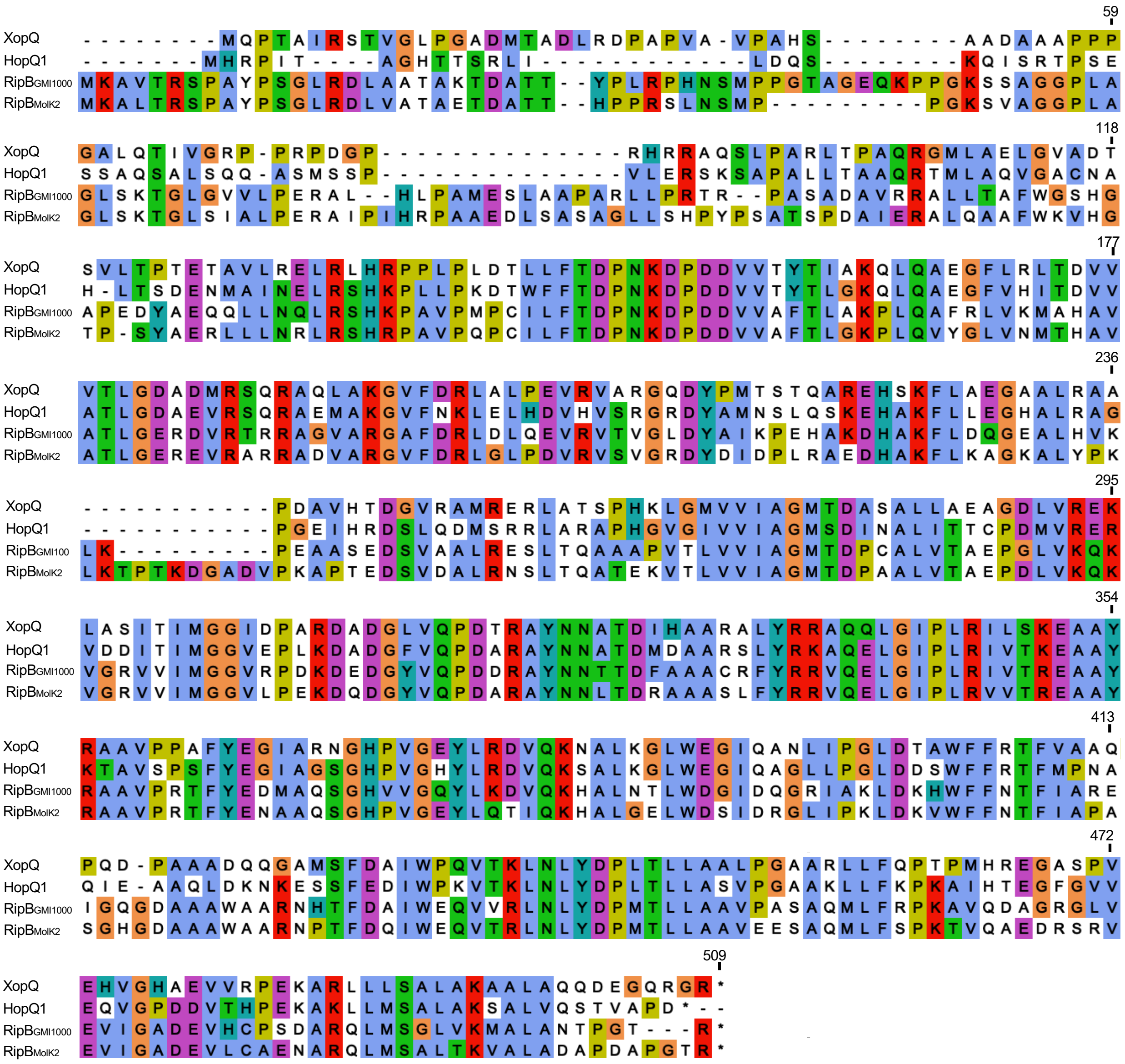


**Supplementary Figure 2.** Protein alignment of XopQ, HopQ1 and RipB. Protein sequences from *Xanthomonas euvesicatoria* 85-10 (XopQ), *Pseudomonas syringae* DC3000 (HopQ1), and the *Ralstonia solanacearum* strains GMI1000 and MolK2 were used to generate the alignment using ClustalO.

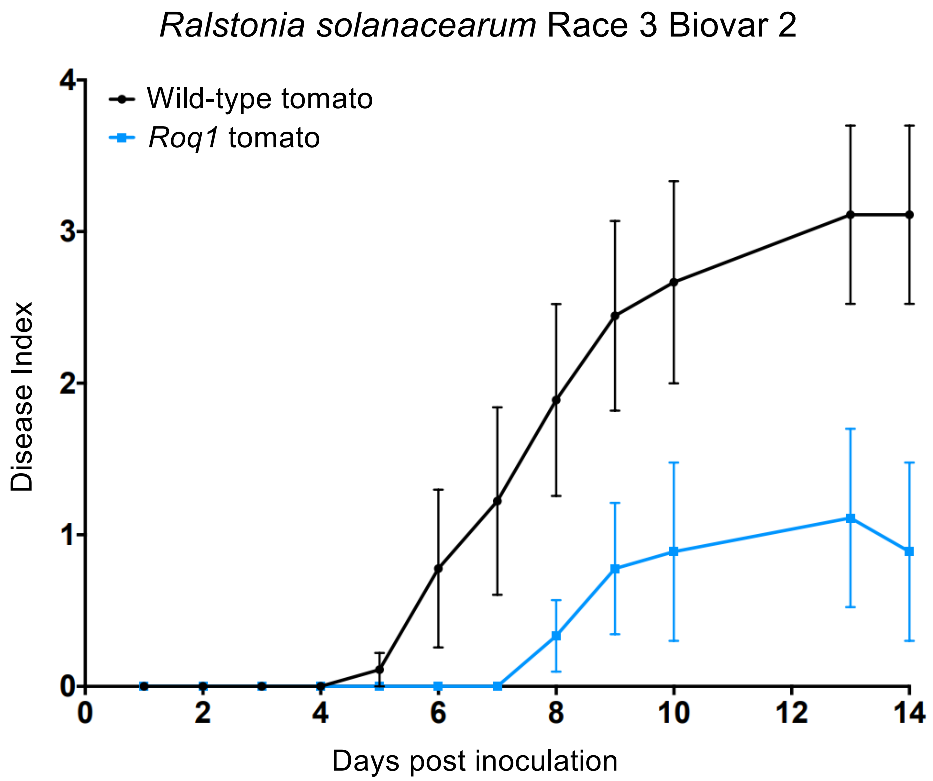


**Supplementary Figure 3.** Disease assay with *Ralstonia solanacearum* Race 3 Biovar 2. Unwounded wild-type tomato plants (cv. Florida 8000) and tomato plants expressing Roq1 were soil-soak inoculated with *R. solanacearum* Race 3 Biovar 2 strain UW551. Disease symptoms were scored over the course of two weeks, with a Disease Index of 0 corresponding to no symptoms and a Disease Index of 4 corresponding to complete wilting. Error bars indicate standard error.

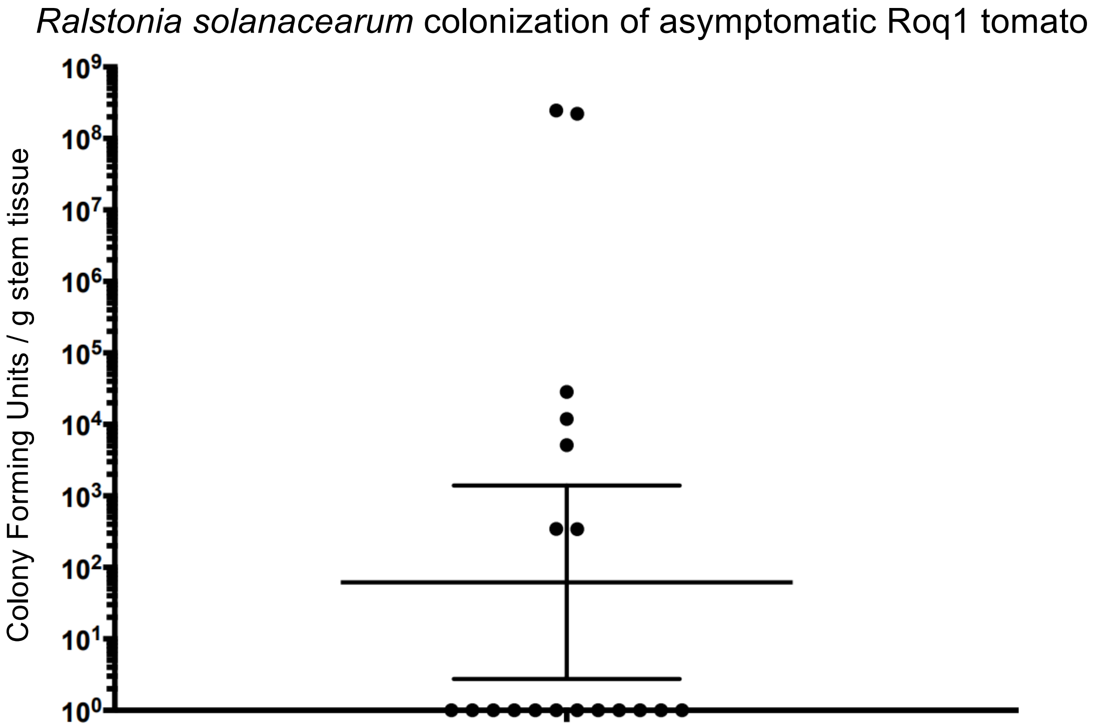


**Supplementary Figure 4.** Colonization of asymptomatic Roq1 tomato plants following inoculation with *Ralstonia solanacearum*. Tomato plants expressing Roq1 were infected using the soil soak method. After two weeks all wild-type tomato plants had wilted but the Roq1 tomato plants appeared healthy with no or minimal disease symptoms. Bacterial population sizes in asymptomatic plants were measured by homogenizing and serial dilution plating mid-stem sections to determine colony forming units per gram stem. Out of 19 plants tested, seven had low or moderate colonization. No *R. solanacearum* cells were detected in stem tissue from the other twelve plants (limit of detection = 100 CFU/gm).
